# Experimental evolution under biased sex ratios: phenotypic and genomic responses in the bulb mite, *Rhizoglyphus robini*

**DOI:** 10.64898/2026.01.23.701246

**Authors:** Sebastian Chmielewski, Jonathan M. Parrett, Mateusz Konczal, Agnieszka Szubert-Kruszyńska, Aleksandra Łukasiewicz, Jacek Radwan

**Affiliations:** Evolutionary Biology Group, Adam Mickiewicz University, ul. Uniwersytetu Poznańskiego 6, 61-614 Poznań, Poland; Institute of Human Biology and Evolution, Adam Mickiewicz University, ul. Uniwersytetu Poznańskiego 6, 61-614 Poznań, Poland

**Keywords:** sexual selection, sex ratio, sexual conflict, sexual antagonism, male harm, experimental evolution

## Abstract

Sexual selection may increase population fitness by favouring high-condition individuals and accelerating the purging of deleterious alleles. However, it can also reduce population fitness through intra- and interlocus sexual conflict by promoting male-benefit traits that harm females and maintain polymorphism at sexually antagonistic loci. The balance between these opposing forces remains unresolved, yet it has major consequences for how sexual selection shapes population fitness and genome-wide variation. To explore the genomic and phenotypic effects of sexual selection and sexual conflict, we evolved replicated bulb mite (Rhizoglyphus robini) lines for 28 generations under male- versus female-biased sex ratios and combined phenotypic assays with whole-genome resequencing. Female fecundity and inbreeding depression did not differ between treatments, and genomic analyses revealed no treatment effect on the loss of rare, putatively deleterious SNPs. Contrary to expectations, males from male-biased lines were less harmful to stock females than males from female-biased lines. Genome-wide nucleotide diversity declined similarly across generations in both treatments, although synonymous exonic diversity declined more slowly in male-biased lines. While only a few SNPs diverged consistently between treatments, we identified large treatment-specific haplotype blocks indicating that multiple genomic regions were involved in response to sex-ratio manipulation. Overall, our results indicate that sex ratio manipulation drives evolution of male harm to females and widespread haplotype frequency changes without clear evidence for enhanced purging or maintenance of genetic diversity. The response thus appears to reflect adaptation to altered level of reproductive competition, but without measurable consequences for population fitness and genetic diversity.

**Significance statement:** Sexual selection is often proposed to improve population fitness by removing deleterious mutations, yet it can also favour traits that harm the opposite sex; consequently, it remains unclear whether stronger reproductive competition reliably enhances population viability. By evolving bulb mite populations under strongly male- or female-biased sex ratios, we found that male-biased populations did not purge genetic load more effectively, while the genomic response to sex-ratio bias was highly polygenic. In contrast to our predictions, males from male-biased lines were less harmful to females than males from female-biased lines. Overall, our results show that sex-ratio bias can reshape male phenotypes and generate patterns of genomic divergence, but without any significant effect on population fitness.

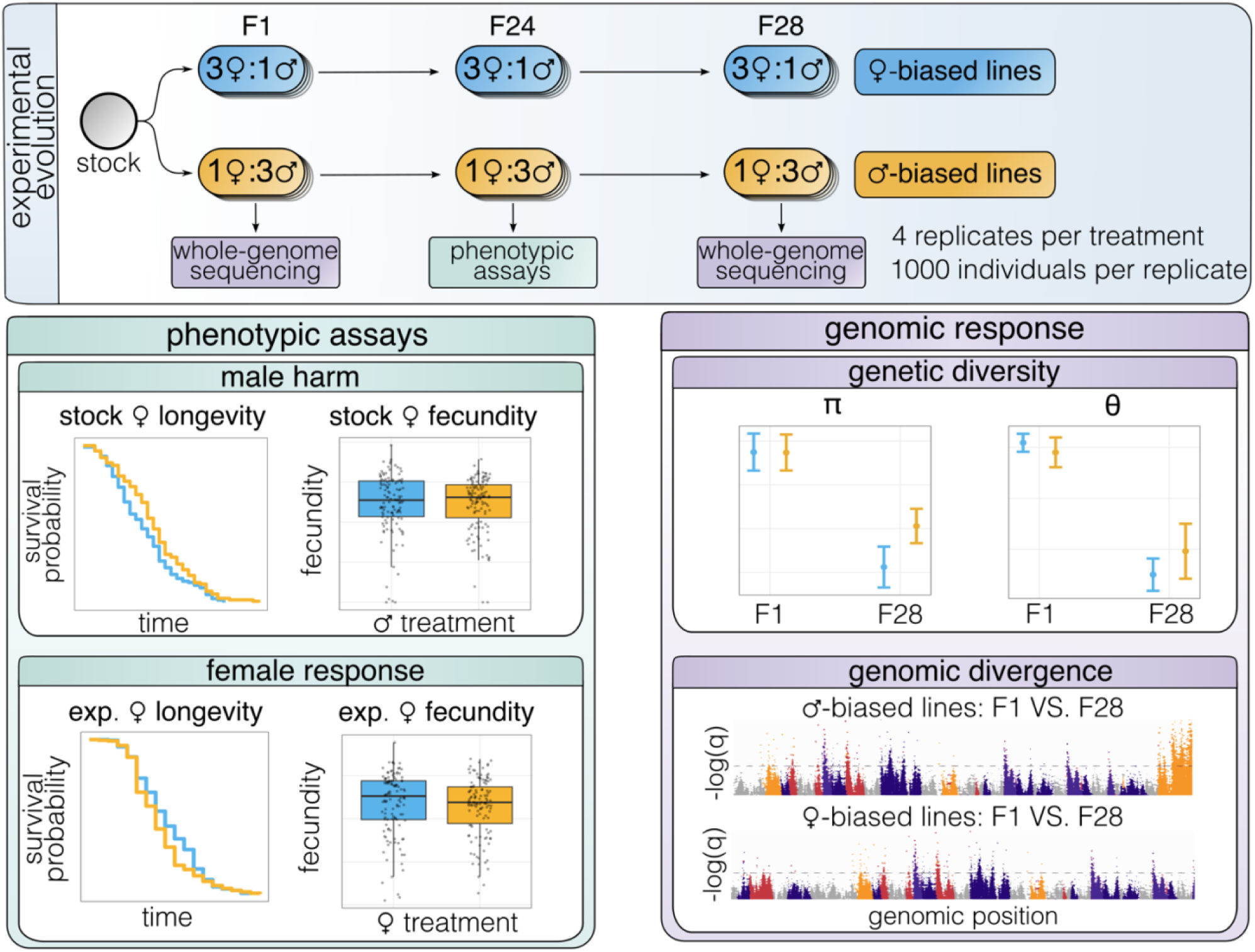

## 1. Introduction

Sexual selection, a process arising from reproductive competition, leads to the evolution of spectacular traits involved in mate choice or competition for mating and fertilisation (Andersson, 1994; Birkhead & Pizzari 2002). Because sexual selection can enhance or oppose other modes of selection, it has a potential to shape genetic variance in populations (Parrett et al., 2022; Rowe & Houle, 1996), with consequences for key evolutionary processes such as adaptation (Lorch et al., 2003; Whitlock & Agrawal, 2009), speciation (Janicke et al., 2018; Ritchie, 2007) and extinction (Kokko & Brooks, 2003; Martínez-Ruiz & Knell, 2017; Morrow & Fricke, 2004).

Sexual selection has the potential to promote beneficial alleles under environmental change (Iglesias-Carrasco et al., 2024; Lorch et al., 2003; Parrett & Knell, 2018) or enhance removal of deleterious alleles from populations (Whitlock & Agrawal, 2009). This is because success in reproductive competition depends on the individual’s general health and vigour (often referred to as condition; Rowe & Houle, 1996), which is negatively affected by maladaptive variants genome-wide (Chen et al., 2025; Dugand et al., 2019; Syed et al., 2025). In consequence, sexual selection can improve population fitness by increasing the strength of purifying selection and thereby reducing deleterious load (Cally et al., 2019; Grieshop et al., 2021), or counteracting extinction risk during population bottlenecks (Jarzebowska & Radwan, 2010; Lumley et al., 2015).

Furthermore, sexual selection may shape adaptive potential of populations by maintaining genetic variance (Barson et al., 2015; Johnston et al., 2013; Ruzicka et al., 2019). Costly sexually selected traits are likely to be involved in fitness trade-offs (Radwan et al., 2016), and the resulting negative pleiotropy, while reducing any population benefits of sexual selection mentioned above, may promote maintenance of genetic variance in populations (Zajitschek & Connallon, 2018). One widespread source of such negative pleiotropy arises from sexually antagonistic selection, i.e., selection on alleles that increase fitness in one sex but decrease it in the other (Bonduriansky & Chenoweth, 2009; Chippindale et al., 2001; Ruzicka et al., 2019; Sayadi et al., 2019). Trade-offs may also involve negative pleiotropic effects of sexually selected traits on other fitness components (Johnston et al., 2013). Moreover, sexual selection can reduce adaptive genetic variation due to lowered effective population size associated with increased variance in male reproductive success (Borgia, 1979; Parrett et al., 2022).

Furthermore, sexual selection can also affect the rate of evolutionary divergence among populations. Divergence in mate preferences can drive divergence in sexual ornaments and underlying genetic variants (Servedio, 2007). Another route to divergence is inter-locus sexual conflict. Because male reproductive success is often strongly dependent on fertilisation success (Janicke et al., 2016; Lehtonen & Parker, 2024), adaptations that increase male success in competition for mates or in sperm competition can be favoured even when they impose costs on females (Arnqvist & Rowe, 2005; Parker, 2006). For example, males may enforce mating even if it increases predation risk for their mates (Arnqvist, 1989) or manipulate female physiology to reduce remating even if it decreases female lifespan (Chapman et al., 1995). Such conflict can lead to various types of dynamics of male adaptation and female counter-adaptation, as increased resistance or choosiness (termed as sexually antagonistic coevolution). Such dynamics under some scenarios may contribute to divergence among populations and ultimately to reproductive isolation (Gavrilets & Waxman, 2002; Hayashi et al., 2007).

Because the strength of sexual selection is likely to be correlated with the strength of sexual conflict (Gómez-Llano et al., 2024), the net effect of reproductive competition on genome-wide genetic variation mentioned above is difficult to predict. For example, both sexually antagonistic selection and inter-locus sexual conflict can erase any adaptive benefits of sexual selection arising from more efficient purifying selection- the former by favouring alleles that improve male fitness while decreasing female fitness (Chippindale et al., 2001; Harano et al., 2010), and the latter by weakening natural selection if high-condition females experience disproportionate harassment or harm (Chenoweth et al., 2015; Long et al., 2009). Empirical data are therefore needed to understand how sexual selection and sexual conflict shape genetic variance relevant to fundamental evolutionary processes of adaptation, speciation and extinction, which is now facilitated by increasing accessibility of genomic resources to non-model organisms (Parrett et al., 2022; Sayadi et al., 2019; Syed et al., 2025). A powerful approach to study effects of these processes on genome-wide variation is experimental evolution followed by re-sequencing, which allows genetic changes to accumulate over generations of selection while controlling for any confounding variables (Kawecki et al., 2012).

Here, we use experimental evolution in the bulb mite *Rhizoglyphus robini* (Acari, Acaridae) to understand how sexual selection and conflict shape genome-wide patterns of variation. Bulb mites are a useful model of sexual selection and sexual conflict, with males showing sexually selected, heritable dimorphism: fighter males bearing weapon (enlarged third pair of legs) used in lethal fights, whereas scrambler males are unarmed and non-aggressive (Parrett et al., 2023; Radwan et al., 2000). While the weapons help males to increase fitness (Radwan & Klimas, 2001), genetic variants associated with weapon expression appear to have detrimental effects on female fitness (Łukasiewicz et al., 2020; Plesnar-Bielak et al., 2014; but see Parrett et al., 2023). Furthermore, the mating system generates strong sperm competition (Radwan, 1997), which selects for high mating rates in males, harmful to their female partners (Kołodziejczyk & Radwan, 2003).

Parrett et al. (2022) artificially selected for or against male weapons to test how sexual selection associated with male aggression shapes genetic variance. Resequencing revealed that populations selected for fighters carried a lower load of deleterious mutations compared to scrambler populations. At the same time, there was little evidence for balancing selection in fighter populations. However, the experiment of Parrett et al. (2022) did not capture important aspects of sexual selection indirectly related to male-male aggression, as selection on morph frequencies does not manipulate the intensity of sperm competition and associated inter-locus sexual conflict (Radwan, 1997). Here, to address this, we experimentally evolved replicated populations of *R. robini* by manipulating sex ratios for 28 generations. Skewing sex ratio toward males is expected to increase selection on traits associated with sperm competition, mating rates and female harassment by males. Thus, male-biased sex ratio is expected to increase the intensity of sexual selection and conflict (Janicke & Morrow, 2018; Kvarnemo & Ahnesjo, 1996; McDonald et al., 2025). However, if increased harassment selects females to reduce resistance or choosiness, precopulatory sexual selection on males may be relaxed (Arnqvist & Rowe, 2005; Fitze & Le Galliard, 2008). In *R. robini*, increased competition for mates influences several levels of sexual selection known to be operating in this species, including probability of female remating which mediates sperm competition and sexual conflict intensity (Radwan & Siva-Jothy, 1996; Kolodziejczyk and Radwan 2003).

After 24 generations of experimental evolution, we tested whether sex-ratio manipulation affected female fitness and male harm to females. If higher reproductive competition among males enhances selection against unconditionally deleterious variants (i.e. with negative effect on fitness of both sexes), we expect females from male-biased lines to show higher fecundity than females from female-biased lines. This difference should be more pronounced under inbreeding, which reveals effects of (partially) recessive mutations (Łukasiewicz et al., 2020; Lumley et al., 2015). Conversely, if increased reproductive competition among males enhances selection for sexually antagonistic alleles, we expect reduced female fitness in male-biased line (Chippindale et al., 2001). If male-biased sex ratios lead to enhanced inter-locus sexual conflict by favouring adaptation in males which harm their mating partners, males evolving under male-biased ratios are expected to impose greater harm on females than males evolving under female-biased sex ratio (Le Galliard et al., 2005). Because increased male harm can select for female counter-adaptations that restore fitness (Rice, 1996; Tilszer et al., 2006), females exposed to harmful males have comparable fitness to females that evolved under lower male harm. Therefore, we also compared fitness of females evolved under male- and female-biased sex ratios after exposure to ancestral (stock) males known to be harmful to females (Kołodziejczyk & Radwan, 2003).

In addition to phenotypic assays, we pool-sequenced our experimental evolution lines after 28 generations and tested the following predictions. If increased male-male competition enhances selection against unconditionally deleterious variants, genetic variance should decline more strongly in male-biased lines compared to female-biased lines. Because deleterious mutations are expected to segregate at low frequencies, the difference should be particularly pronounced in Watterson’s estimator (hereafter θ, Watterson, 1975), a measure of diversity based on the number of segregating sites and thus sensitive to the loss of rare variants. If, instead, sexual selection is associated with widespread sexual conflict maintaining sexually antagonistic variants at intermediate frequencies, the effect of manipulation of sexual selection should mostly affect nucleotide diversity π, and to do so in a less predictable way. This is because π has a maximum value at even proportions of alleles, so how π will change will depend on initial equilibrium frequency and in which direction the equilibrium would move following experimental evolution. Finally, it is possible that sexual selection and/or conflict favours genetic variants associated with particular male and female traits involved, leaving localised signals of genomic divergence (Wiberg et al., 2021). Therefore, in addition to testing the effect of sex-ratio manipulation on genome-wide diversity, we characterised nucleotide polymorphism and genomic regions that responded to sex-ratio manipulation.

## 2. Results

### 2.1. Effects of sex-ratio selection on male harm to females

We evolved replicate populations under female-biased (3♀:1♂; n = 4 lines) or male-biased (1♀:3♂; n = 4 lines) sex ratios for 28 generations. To reduce the effect of genetic drift, which can overwhelm the effect of selection (Kofler & Schlötterer, 2014), we used census size of 1,000 adults per line.

To test whether evolution under the male-biased sex ratio enhanced purging of deleterious alleles, we compared inbreeding depression in female fecundity between sex-ratio regimes. There was no significant inbreeding by sex-ratio treatment interaction, indicating that inbreeding affected female fecundity similarly in both regimes (*χ^2^* = 0.83, *df =* 1, *P* = 0.77, Figure 1A). In the final model, which included only the main effects of inbreeding and selection regime, inbred females showed significantly reduced fecundity relative to outbred females (*χ^2^* = 9.82, *df* = 1, *P* = 0.0018), whereas fecundity did not differ between male- and female-biased regimes (*χ^2^* = 0.92, *df* = 1, *P* = 0.33).

**Fig. 1.**
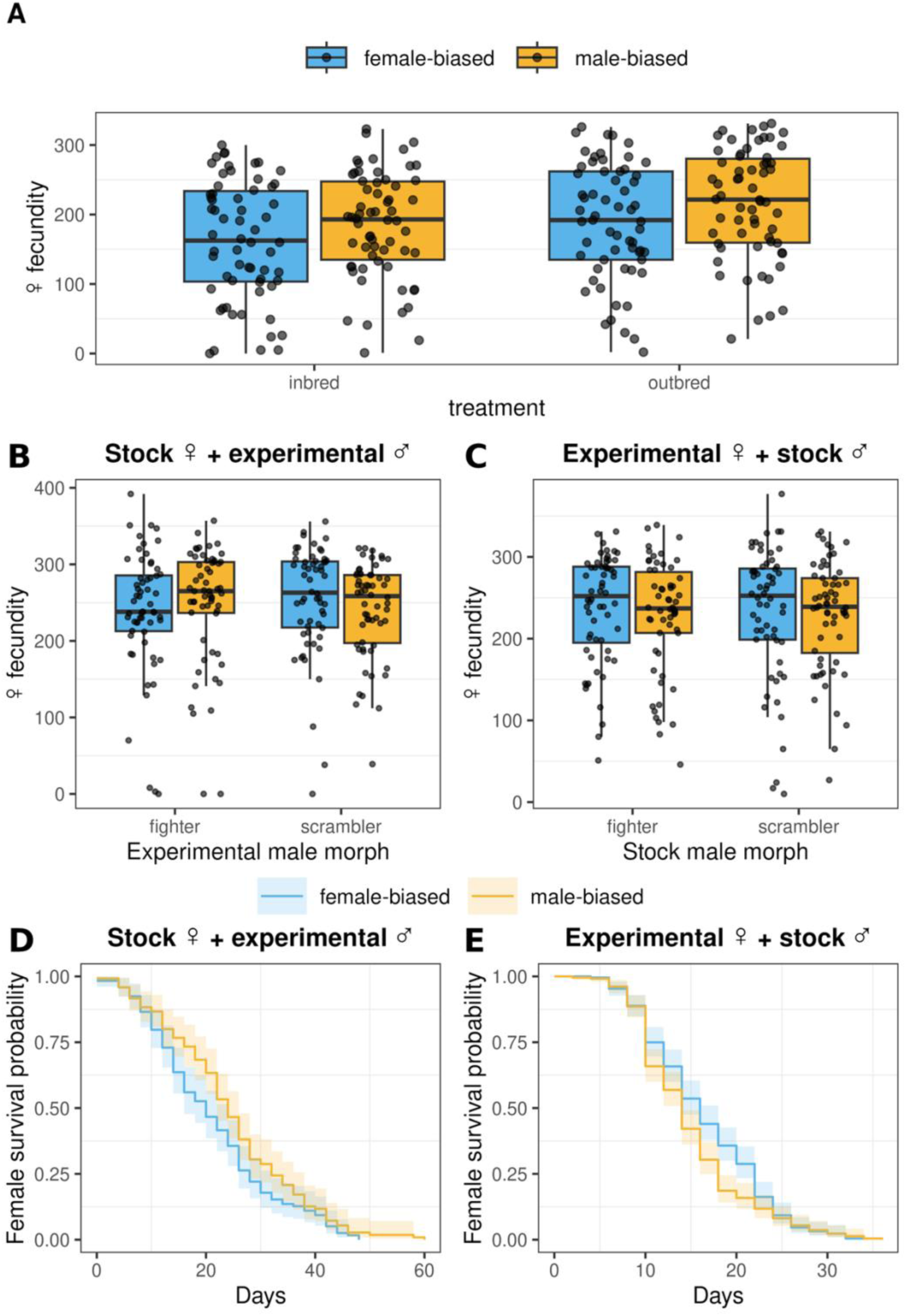
Phenotypic responses to sex-ratio selection in bulb mites. Colours indicate the selection regime of the experimental lines (blue: female-biased; yellow: male-biased). (A) Fecundity of inbred and outbred experimental females from male- and female-biased lines. Inbreeding reduced female fecundity, but fecundity did not differ between sex-ratio treatments. (B) Fecundity of stock females mated with experimental males of either morph (fighter or scrambler) from male- and female-biased lines; fecundity was not affected by selection regime or male morph. (C) Fecundity of experimental females from male- and female-biased lines mated with stock males of either morph; fecundity did not differ with selection regime or male morph. (D) Survival of stock females paired with experimental males; females mated with males from male-biased lines lived longer than those mated with males from female-biased lines. (E) Survival of experimental females mated with stock males; females from male-biased lines tended to have shorter lifespans than females from female-biased lines. Survival curves are Kaplan-Meier estimates and shaded areas represent 95% confidence intervals.

We then tested predictions of sexual conflict, i.e. whether higher reproductive competition leads to evolution of increased male harm to their female partners, resulting in counter-adaptation restoring female fitness. To test whether males that evolved under different sex ratios differed in their effects on female fitness, we paired stock females with males from the experimental lines and checked female longevity and fecundity. Stock females paired with males from male-biased lines lived longer than those mated with males from female-biased lines (*χ^2^* = 4.21, *df* = 1, *P* = 0.04; Figure 1D). Female longevity was not affected by male morph (*χ^2^* = 0.05, *df* = 1, *P* = 0.83). Stock female fecundity did not differ between sex-ratio regimes (*χ^2^* = 0.03, *df* = 1, *P* = 0.87, Figure 1B) and was not affected by male morph (*χ^2^*= 0.25, *df* = 1, *P* = 0.61).

Finally, to assess whether females evolved in response to sex-ratio manipulation, we paired experimental females with stock males and measured female longevity and fecundity. Females from male-biased lines tended to have shorter lifespans than females from female-biased lines, although this effect was marginally non-significant (*χ^2^* = 0.08, *df* = 1, *P* = 0.059; Figure 1E). Female fecundity did not differ between sex-ratio regimes (*χ^2^* = 0.16, *df* = 1, *P* = 0.69; Figure 1C) and was not affected by the morph of the stock male (t = 0.07, *df* = 1, *P* = 0.79).

### 2.2. Autosomal genetic diversity

To quantify genomic responses to experimental evolution, we performed whole-genome pool-sequencing of ancestral (generation 1) and evolved (generation 28) samples from each replicate line using pools of 100 females and 100 males per line and generation. Using the PoPoolation2 pipeline (Kofler, Pandey, et al., 2011), we identified 6.13 million high-quality SNPs across the genome, including 702,656 SNPs on the X chromosome. All diversity analyses reported here were restricted to autosomal SNPs.

To assess the genetic diversity, we calculated nucleotide diversity (π) and Watterson estimator (θ). These measures were calculated within four SNP classes: whole exons, non-synonymous sites, synonymous sites, and 10kb windows, which comprised both exons and intergenic regions. We observed a significant decline in π and θ between the ancestral and evolved populations in all analysed SNP classes (Figure 2; Table 1). Within exons and synonymous sites, male-biased lines showed significantly lower decline in π compared to female-biased lines (Figure 2), as indicated by a significant treatment by generation interaction (Table 1), but the interaction was not significant for 10kb windows and non-synonymous sites. Sex-ratio treatment did not significantly affect the change in θ between treatments over generations.

**Fig. 2.**
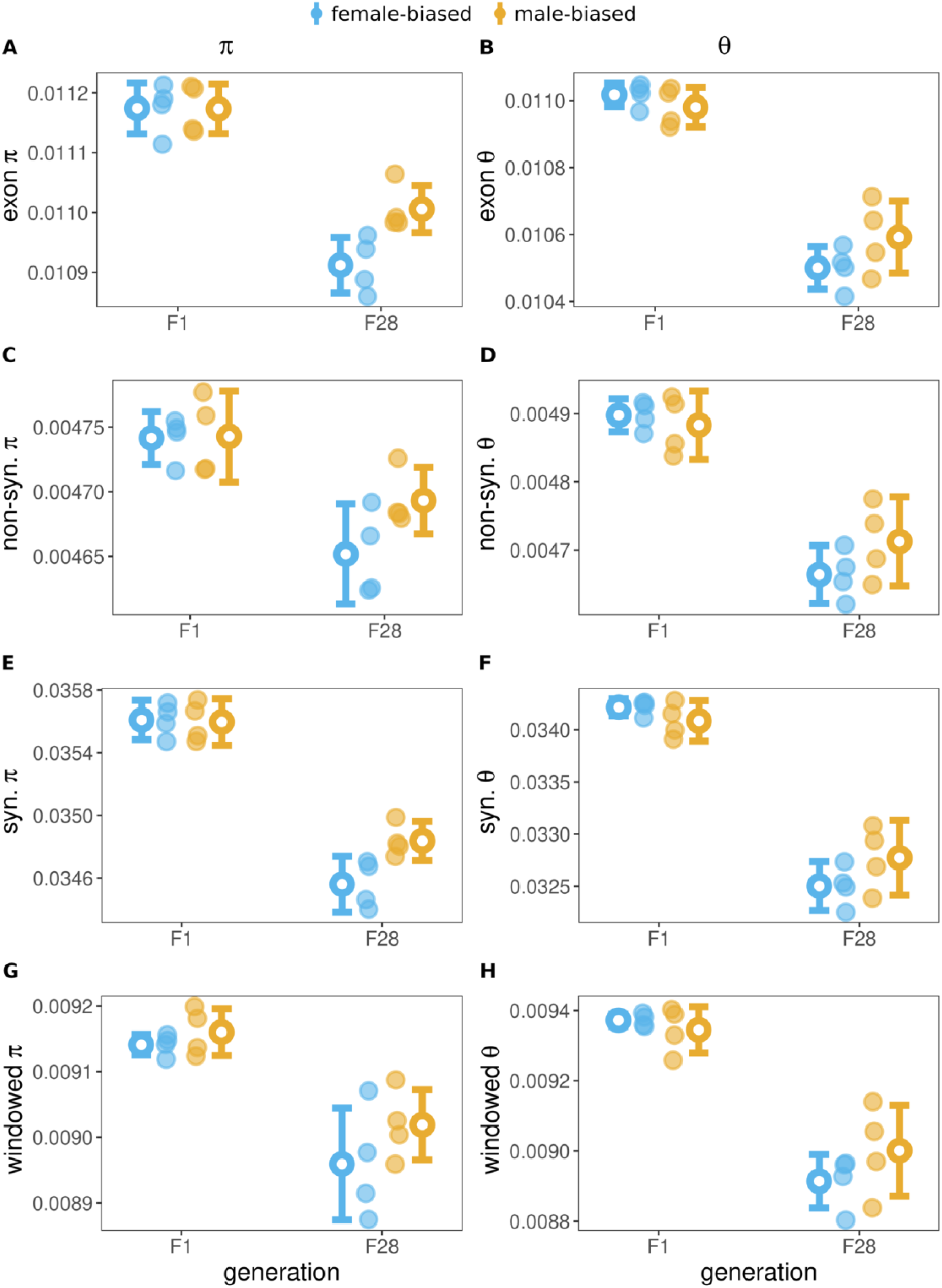
Changes in autosomal diversity patterns. Left panels show nucleotide diversity (π); right panels show Watterson’s estimator (θ). Faded points represent line means; open points and whiskers denote treatment means ± 1 s.d. across lines.

**Table 1.** Results of analysis of variance of genetic diversity measures. *P* values below 0.05 are in bold.

| measure | class | term | $F_{(1,12)}$ | *P* |
| --- | --- | --- | --- | --- |
| $\pi$ | exon | treatment | 4.785 | **0.0492** |
|  |  | generation | 102.747 | **<0.0001** |
|  |  | treatment:generation | 4.989 | **0.045** |
|  | syn | treatment | 4.527 | **0.055** |
|  |  | generation | 212.515 | **<0.0001** |
|  |  | treatment:generation | 5.442 | **0.038** |
|  | nonSyn | treatment | 2.643 | 0.129977 |
|  |  | generation | 28.127 | **<0.0001** |
|  |  | treatment:generation | 2.340 | 0.152 |
|  | 10kb windows | treatment | 2.127 | 0.170 |
|  |  | generation | 35.759 | **<0.0001** |
|  |  | treatment:generation | 0.563 | 0.467 |
| **θ** | exon | treatment | 0.584 | 0.459 |
|  |  | generation | 161.616 | **<0.0001** |
|  |  | treatment:generation | 3.302 | 0.094 |
|  | syn | treatment | 0.475 | 0.504 |
|  |  | generation | 223.709 | **<0.0001** |
|  |  | treatment:generation | 3.952 | 0.07 |
|  | nonSyn | treatment | 0.705 | 0.42 |
|  |  | generation | 98.386 | **<0.0001** |
|  |  | treatment:generation | 2.393 | 0.148 |
|  | 10kb<br>windows | treatment | 0.534 | 0.479 |
|  |  | generation | 95.323 | **<0.0001** |
|  |  | treatment:generation | 1.931 | 0.19 |

Both genetic drift and purifying selection are expected to contribute to the observed decrease in π and θ. To separate these two processes, we compared changes between generations in synonymous sites (which mostly indicate genetic drift) and in non-synonymous sites (reflecting selection), following Parrett et al. (2022; see *Methods*). Values of Δπ or Δθ close to 0 indicate that nonsynonymous diversity changed to a similar extent as synonymous diversity, consistent with drift alone. Negative values (Δπ or Δθ < 0) indicate an excess loss of nonsynonymous variation relative to drift expectations (purifying selection), whereas positive values (Δπ or Δθ > 0) indicate maintenance or increase of non-synonymous diversity relative to drift.

For Δπ, 95% confidence intervals of line-averaged values overlapped with zero in both treatments (Figure 3A, 95% CI female-biased: −0.005 to 0.003; 95% CI male-biased: - 0.003 to 0.005), and Δπ did not differ significantly between male- and female-biased lines (t-test, t = −1.06, *df* = 6, *P* = 0.33). This suggests that changes in nucleotide diversity were mostly driven by genetic drift. In contrast, Δθ was significantly negative in both treatments (95% CI female-biased: −0.014 to −0.005, 95% CI male-biased: −0.009 to −0.001, Figure 3B), indicating that the number of segregating sites declined faster than expected under drift alone, consistent with purifying selection. However, Δθ did not differ significantly between the treatments (t-test, t = −1.86, *df* = 6, *P* = 0.11).

**Fig. 3.**
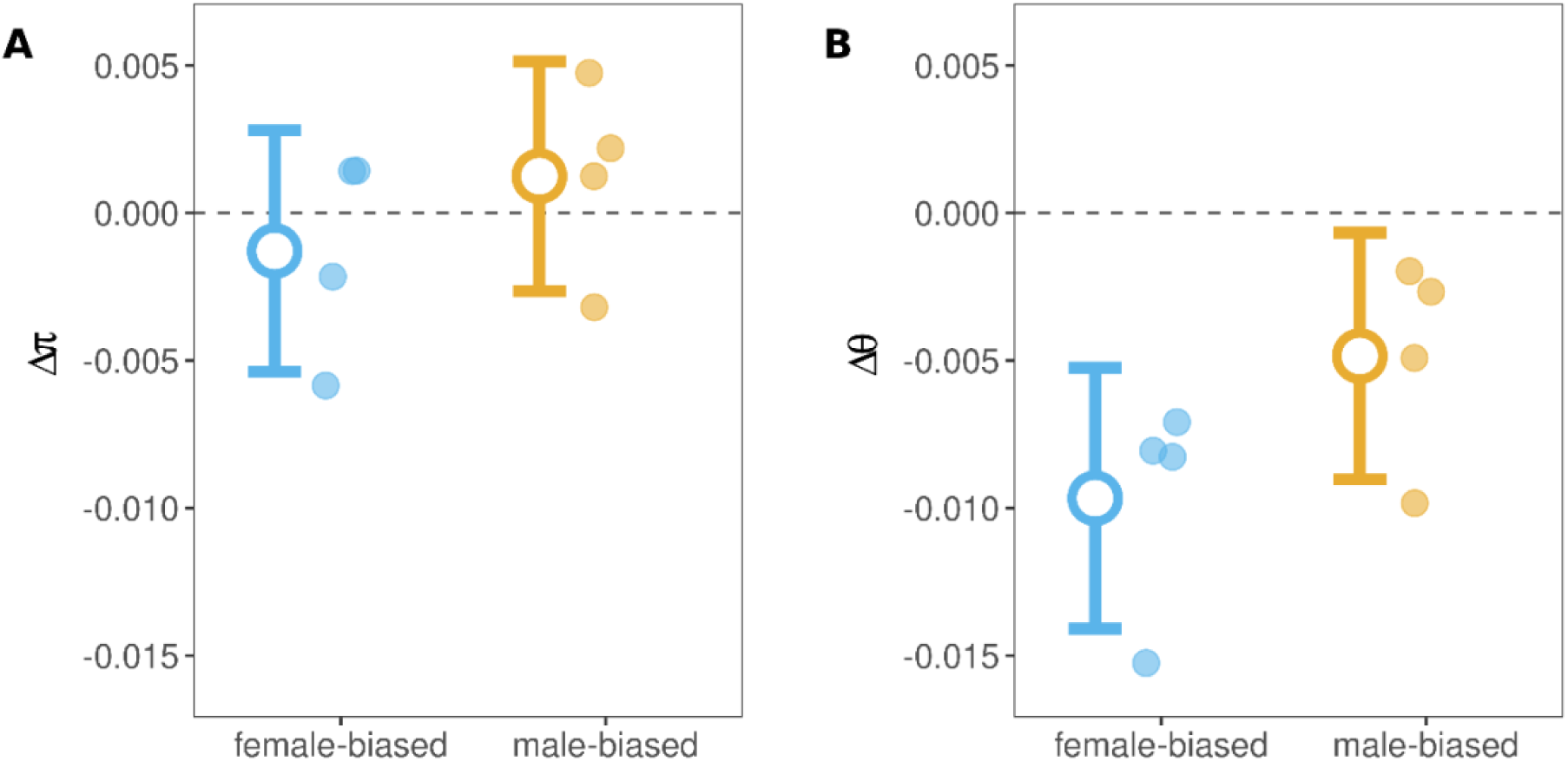
Relative change in non-synonymous diversity after controlling for genome-wide synonymous change indicated by Δπ (A) and Δθ (B). Faded points show values for individual lines; open points and whiskers indicate treatment means and 95% confidence intervals for female-biased (blue) and male-biased (yellow) lines.

Estimates of effective population size based on allele-frequency changes between ancestral and evolved samples did not differ significantly between the sex-ratio regimes (Figure S1, t = 0.394, *df* = 6, *P* = 0.71; mean of male-biased lines = 397, range: 356 - 478; mean of female-biased lines = 409, range: 395 - 441). Estimates of effective population sizes for all lines are provided in Table S2.

### 2.3. Divergence between basal and evolved lines

To identify SNPs that diverged during experimental evolution, we applied the ACER test, which detects consistent allele-frequency changes across replicate lines while accounting for genetic drift and sampling error in pool-seq data (Spitzer et al., 2020). We first focused on SNPs which frequency changed significantly in both sex-ratio regimes. Among 17,918 such SNPs, only five showed allele-frequency shifts in opposite directions between female- and male-biased lines (Figure 4, Table S3). Of these five SNPs, one was located within a protein-coding gene annotated as a homologue of a nuclear hormone receptor HR96 (SNP position chr8: 6,268,648). Three SNPs were located within 673 bp upstream of the nearest genes (two near a gene encoding a tripartite motif-containing protein and one near a gene of unknown function), and the remaining SNP was located 719 bp upstream of a heat-shock protein gene.

**Fig. 4.**
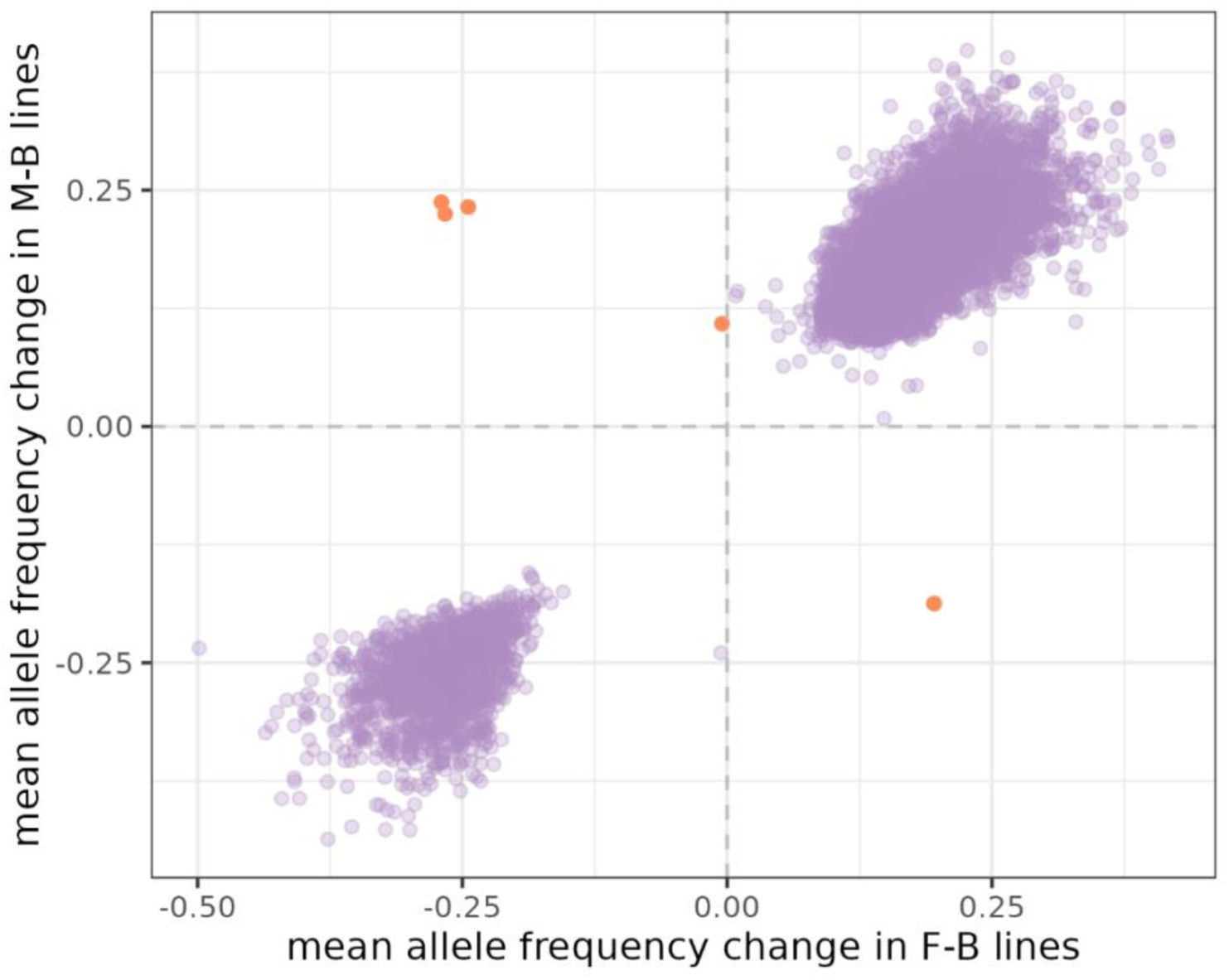
Correlation of allele frequency change between female- and male-biased lines. Only SNPs that significantly changed their frequency between basal and evolved populations in both treatments, as indicated by the ACER test, are shown. Violet points indicate concordant allele frequency changes between treatments, while orange points indicate SNPs that changed their frequency in opposite directions in male- and female-biased lines.

For the remaining 17,913 SNPs that changed significantly in both treatments, allele-frequency changes were highly positively correlated between male- and female-biased lines (Figure 4, Pearson’s r = 0.96, *P* < 0.0001), indicating that most of these loci likely contribute to adaptation to the common experimental conditions rather than to divergence driven by sex-ratio manipulation.

Using the ACER test, we also identified SNPs which frequencies shifted significantly in only one sex-ratio treatment. In female-biased lines, 49,228 SNPs changed in frequency with no detectable response in male-biased lines. Based on shared allele-frequency trajectories, these SNPs clustered into 46 independent haplotype blocks (Figure 5A). In male-biased lines, 79,390 SNPs changed frequency exclusively in that treatment, segregating in 58 haplotype blocks (Figure 5B).

**Fig. 5.**
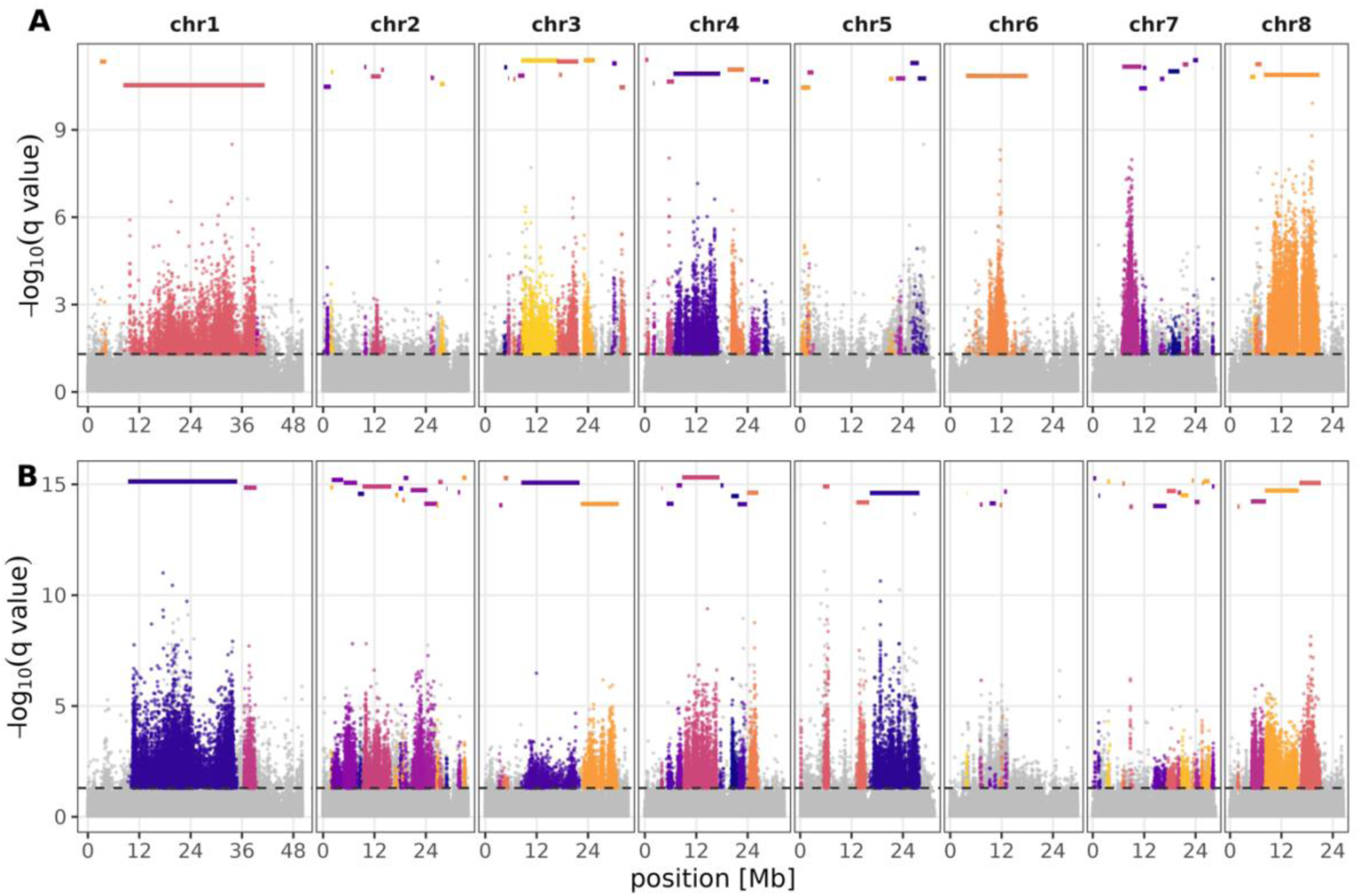
Haplotype blocks identified from treatment-specific SNPs. (A) Haplotypes inferred from SNPs that diverged exclusively in female-biased lines. (B) Haplotypes inferred from SNPs that diverged exclusively in male-biased lines. Individual haplotypes are color-coded. The x-axis shows genomic position; the y-axis shows the negative log-transformed q-values from the ACER test. The dashed horizontal line indicates q = 0.05.

Within each treatment, we performed functional analysis of selection targets. Because selected haplotype blocks in E&R experiments carry many linked variants that hitchhike with loci under selection, we restricted our analysis to the top 10% of significant SNPs with the lowest q-values within each haplotype, which we refer to as selection targets. In female-biased lines, selection targets were enriched for membrane-associated functions (GO:0016020; Table 2). In male-biased lines, selection targets were enriched for transcription coregulation (GO:0003712) and gene regulation via histone acetylation (GO:0004402). Haplotype length distributions were similar between treatments (Wilcoxon rank-sum test: W = 1391, *P* = 0.47) and ranged from 2.2 kb to 33 Mb (Figure S4).

**Table 2.** GO term enrichment for selection targets (top 10% of significant SNPs per haplotype) that diverged exclusively in female- or male-biased lines.

| treatment | GO term | GO description | adjusted $P$ | gene ratio |
| --- | --- | --- | --- | --- |
| female-biased | 0016020 | membrane | 0.011 | 0.1 |
| male-biased | 0003712 | transcription coregulator activity | $< 0.0001$ | 0.02 |
| male-biased | 0004402 | histone acetyltransferase activity | 0.002 | 0.02 |

Finally, to facilitate direct comparison with our previous evolve-and-resequence experiment in which sexual selection was manipulated by altering the frequency of alternative male morphs (Parrett et al. 2022), we also used generalized linear models to test for SNPs that diverged consistently between male- and female-biased lines at generation 28. After controlling for false discovery rate (q < 0.05), only a single SNP exceeded the significance threshold. This SNP, located at chr1: 4,172,390, was not situated near any annotated protein-coding gene, and had mean allele frequencies of 0.63 in female-biased lines and 0.84 in male-biased lines (Figure S5)

## 3. Discussion

In the present study, we manipulated intrasexual reproductive competition by evolving replicate populations under male-biased or female-biased sex ratios for 28 generations and tested whether this led to enhanced selection and consequently to increased fitness. We also tested whether increased reproductive competition leads to increased male harm to females, as expected under intersexual conflict theory. Finally, we identified genetic variants which frequencies changed in response to sex-ratio manipulation in order to characterise the genomic basis of the response.

### 3.1. Sex-ratio manipulation did not enhance purging of deleterious variation

At the phenotypic level, we found no support for the prediction that enhanced reproductive competition among males improves population fitness and reduces inbreeding load, the latter often used as a proxy for the load of partially recessive deleterious mutations (Łukasiewicz et al., 2020; Lumley et al., 2015). Our results on genome-wide diversity were consistent with this finding. The only significant effect of sex-ratio manipulation was significantly slower decline in synonymous π across generations in male-biased lines. This pattern does not indicate enhanced purifying selection in male-biased lines, because treatment did not influence the relative change in non-synonymous diversity once genome-wide synonymous change was accounted for, and purifying selection inferred from Δθ was not detectably stronger in male-biased lines. Moreover, estimates of effective population size (Ne) did not differ significantly between sex-ratio treatments, which argues against a simple explanation in terms of stronger drift in one treatment.

Both phenotypic and genome-wide effects of sex-ratio manipulation contrast with previous work in the same system, which used the same starting population but manipulated sexual selection by artificially selecting for distinct male morphs (Parrett et al., 2022). That study increased the proportion of males bearing a condition-dependent weapon in the population, and the selection was highly effective in driving morph frequencies toward fixation. This in turn changed the nature of sexual selection, with the opportunity for lethal fights eliminated in scrambler-fixed selection lines. At the phenotypic level measured after 45 generations, morph proportion manipulation led to higher inbreeding load for female fecundity in scrambler-fixed lines compared with fighter-fixed lines, suggesting that selection on the weapon and/or sexual selection associated with male fights enhanced purging of deleterious recessive mutations. Consistent with these phenotypic effects, scrambler-selected lines had higher frequencies of initially low-frequency, presumably deleterious variants, leading to higher estimates of genetic diversity than in fighter-selected populations, particularly for Watterson’s estimator θ, a parameter particularly sensitive to rare variants. Another striking contrast with our current results is the number of SNPs detected by generalised linear models (GLMs) as significantly diverged between treatments (1 here versus ∼24,000 in Parrett et al. 2022). This paucity of diverged SNPs again suggests that selection in the present study was much weaker than direct selection on male weapon. Here, we purposefully counteracted morph ratio evolution by maintaining equal proportions of fighters and scramblers in both sex-ratio treatments in order to assess the effects of male reproductive competition on components of sexual selection other than inter-male aggression. Our results suggest that these other components either constitute a much weaker force acting on genetic load than that associated with lethal male competition, and/or that biasing sex ratio is less effective in manipulating the strength of sexual selection. Although sex-ratio manipulation is a common approach to manipulate the strength of sexual selection, and has often been reported to increase the opportunity for sexual selection (e.g. House et al., 2019; Souroukis & Cade, 2014; Wacker et al., 2013), some studies do not support such a relationship (Dutta et al., 2022; Head et al., 2008; McDonald et al., 2025). For example, in *Drosophila melanogaster*, a male-biased sex ratio increased sperm competition, but this did not translate into a steeper relationship between male mating success and the offspring number (i.e. the Bateman gradient; McDonald et al., 2025). Regardless of whether male bias universally strengthens sexual selection, our manipulation altered selection on sexually selected traits, as evidenced by evolved differences in male-induced female longevity. Furthermore, another *D. melanogaster* study reported that male-biased ratios generally increase selection against two large effect mutations, also in a context and locus specific manner (Sharda et al., 2024). Consistent with such locus dependence, Arbuthnott and Rundle (2012) found that increasing the opportunity for sexual selection (polygamy vs monogamy) did not accelerate purging of six deleterious alleles.

In addition to differences in the strength of sexual selection, the present study may also differ from Parrett et al. (2022) in selection target, which could be larger in the present study due to the highly polygenic nature of the response associated with multiple aspects of phenotypes being under selection due to sex-ratio manipulation. Highly polygenic adaptation may lead to only small allele frequency changes, and to little overlap between replicate populations, making them difficult to detect (Barghi et al., 2020; Otte et al., 2021). This may explain the contrast between our study and those of Sharda et al. (2024) who studied effect of sex ratio on large-effect mutations.

### 3.2. Evolution of male harm under sex-biased treatments

Despite the absence of detectable treatment effects on fecundity and inbreeding depression, sex-ratio manipulation did affect male-induced harm to females. Stock females paired with males from male-biased lines lived longer than those paired with males from female-biased lines, while female fecundity in these assays did not differ detectably between male treatments. An earlier study in the same species manipulated reproductive competition enforcing monogamy or allowing for polygamous mating and found that eliminating reproductive competition decreased male harm to females, increasing female fecundity and longevity (Tilszer et al., 2006). Furthermore, females from monogamy treatments evolved to be more sensitive to male harm. Similar findings have been reported in *D. melanogaster* when sexual selection (and conflict) were manipulated by varying sex ratios (Nandy et al., 2013, 2014; Wigby & Chapman, 2004). In contrast, we found that males evolving under male biased sex ratios appeared to be less harmful to stock females than males evolving under female biased sex ratio. However, while sexual selection seems typically stronger when sex ratios are more male biased (Janicke & Morrow, 2018; Kvarnemo & Ahnesjo, 1996), this relationship can depend on whether the abundant sex (e.g. males) can monopolise reproductive opportunities; under strongly male-biased sex ratios, monopolisation may be hampered, potentially reducing sexual selection on males (Klug et al., 2010; Kokko et al., 2012). A key point is that sex-ratio bias can alter the relative contribution of pre- versus postcopulatory selection. If male harm is expressed primarily through precopulatory mechanisms, as suggested in bulb mites (Konior et al., 2006; Skwierzyńska & Plesnar-Bielak, 2018), then stronger male bias may shift selection toward postcopulatory traits that enhance sperm competitive ability and fertilisation success, which may be less tightly linked to female harm. This provides a plausible route by which male-biased lines could evolve males that are, on average, less harmful to females in survival assays despite elevated reproductive competition. Consistent with this interpretation, a recent bulb mite study manipulating sex ratio found that more male-biased conditions were associated with a decline in a male-benefit, female-harm allele involved in sexual conflict in this species (Unnikrishnan et al., 2025).

Nevertheless, previous work on the bulb mites showed that sex-ratio manipulation (1:2 vs 2:1 males to females) can generate evolutionary change in traits linked to male reproductive success and male-induced harm to females (Plesnar-Bielak et al., 2020). It therefore remains puzzling why in our experiment, males evolving under male-biased sex ratios appeared to be less harmful to stock females than males from female-biased lines. Possibly, more extreme sex-ratio bias used here, and/or differences in demography and ecology between studies (e.g., population size and density), altered the balance of selection acting on male harm and correlated reproductive traits. While further work is needed to determine why male-biased sex ratio led to the evolution of males that are more benign to females, it is noteworthy that females evolved in a way consistent with relaxed selection caused by male harm: females evolving under decreased male harm (i.e. under female-biased ratios) tended to have higher fecundity when exposed to control males, although this effect was not significant. This mirrors findings (e.g. Nandy et al., 2014; Rice, 1996; Tilszer et al., 2006) that females coevolving with more harmful males can exhibit higher fecundity when exposed to harmful males than females coevolving with more benign males.

### 3.3. Genome-wide response to sex-ratio manipulation

The phenotypic response to sex-ratio manipulation in terms of male effect on female fitness (possibly associated with female resistance evolution), was accompanied by allele-frequency changes across multiple genomic regions. GLMs detected only one SNP that diverged consistently between sex-ratio treatments at generation 28, indicating that large, repeatable treatment differences at frequency of individual loci were rare. In contrast, within-treatment analysis using ACER test identified many SNPs that were significantly selected for in our experimental evolution lines. A substantial proportion of these significant SNPs (12%) was selected consistently in both treatments, likely reflecting adaptation to experimental conditions that differed from those experienced in stock population (e.g. more constrained development time, lower colony density, or food availability).

Only five SNPs were identified as changing in opposite directions between treatments, suggesting limited sexual antagonism associated with our manipulation of the intensity of reproductive competition. Only one of these SNPs was located within a protein-coding sequence, in the homologue of the nuclear hormone receptor HR96. While this gene has known roles in transcriptional regulation and xenobiotic response in other arthropods (Fisk & Thummel, 1995, Ji et al. 2023), the current data do not provide an obvious explanation for divergent selection between sex-ratio treatments.

In contrast to the small number of divergently changing SNPs, many SNPs showed treatment-specific responses (i.e. significant changes in one treatment but not the other). Because selection on a locus can drive many linked SNPs to change in frequency, we therefore summarised genomic responses in terms of haplotype blocks and defined putative targets within those blocks, rather than treating all significant SNPs as independent targets of selection. We identified 46 and 58 haplotypes in female- and male-biased lines, respectively, supporting polygenic response to sex-ratio treatments. Similar genome-wide responses to altered sexual selection have been found in *Drosophila pseudoobscura*, where elevated sexual selection (polyandry) produced stronger genomic signals of selection and a greater number of significant variants than relaxed sexual selection (monogamy; Barata et al., 2023). Moreover, analyses of the same experimental lines showed that genomic divergence was concentrated in discrete ‘islands’, consistent with our observation of clustered, haplotype-level responses (Wiberg et al., 2021). Functional enrichment based on putative selection targets suggested that the most strongly responding SNPs in female-biased lines were enriched in genes associated with membrane-related functions, whereas in male-biased lines they were enriched in categories related to transcriptional coregulation and histone acetylation mechanisms that can have downstream effects via gene regulation (Sterner & Berger, 2000). These categories suggest potentially broad consequences of sex-ratio manipulation. Many haplotypes, including some long ones, did not overlap between treatments, suggesting that female- and male-biased sex ratios favoured different genomic regions. This reinforces conclusions from our phenotypic results suggesting that sex-ratio manipulation can have complex and not easily predictable consequences for selection acting on different traits and their underlying genomic regions, consistent with work in *D. melanogaster* showing locus-specific responses to sex-ratio manipulation (Sharda et al., 2024).

### 3.4. Concluding remarks

Overall, our results show that sex-ratio manipulation can affect the evolution of male traits and drive allele-frequency changes across multiple genomic regions, but these effects are not readily reconciled with standard predictions about how increased reproductive competition should affect the strength and direction of sexual selection. Contrary to expectation that male-biased sex ratio should strengthen sexual selection (Janicke & Morrow, 2018; Kvarnemo & Ahnesjo, 1996) and thereby enhance selection against deleterious mutations (Dugand et al., 2019; Parrett et al., 2022), we did not detect enhanced purifying selection at either the phenotypic or genomic level. Moreover, and again contrary to expectation, enhanced reproductive competition between males did not lead to increased male harm to females, but the opposite pattern emerged. These results highlight that sex-ratio manipulation may have unexpected impact on the intensity of sexual selection, as already noted by other researchers (Kokko et al., 2012; McDonald et al., 2025; Sharda et al., 2024). Yet, despite generally weak and unexpected phenotypic response, sex-ratio manipulation did lead to widespread selection on many genomic regions. The mechanisms underlying these unexpected phenotypic effects, and the extensive genomic evolution that accompanied them, require further investigation.

## 4. Materials & Methods

### 4.1. Experimental evolution

We used the same stock population of *R. robini* as in Parrett et al. (2022). It was established by mixing three outbred populations (Mosina, Kwiejce, and Kraków) to increase genetic diversity, followed by approximately 30 generations of free mating to break linkage disequilibria that could have arisen due to mixing. The experimental design consisted of eight lines of 1000 individuals each: 4 female-biased lines with a 3:1 female-to-male ratio and 4 male-biased lines with a 1:3 female-to-male ratio. All other experimental procedures followed Parrett et al. (2022).

Briefly, the mite colonies were reared in tightly sealed plastic dishes (9 x 7 x 4.5 cm, length x width x height) with a 1 cm plaster-of-Paris base. To allow for airflow, the lid had small holes that prevented mites from escaping. High humidity was ensured by soaking a plaster-of-Paris base with water. Mites were fed with dry yeast provided *ad libitum*.

At each generation, upon reaching maturity, the desired number of random males and females were manually selected and transferred to new dishes. To separate the effects of morph competition, previously studied in Parrett et al. (2022), from the overall effects of sexual selection, we controlled the morph ratio by selecting equal numbers of fighter and scrambler males in both sex-ratio treatments. After selection, mites were allowed to interact for six days and eggs were separated from adults by rinsing the colony over a fine mesh that allowed eggs to pass through while retaining adult mites. Adults were then transferred to new dishes to lay eggs for one additional day. These eggs were used to found next generation, and adult mites were discarded. Frequent mating (several copulations a day, Radwan & Siva-Jothy, 1996), coupled with last sperm precedence observed in bulb mites (Radwan, 1997), implies that the eggs used to start next generation were fertilized mostly with the sperm of the selected males available during the interaction period. Because there were three times as many ovipositing females in female-biased treatment, we evenly mechanically removed approximately two-thirds of the dish surface area in the female-biased lines after the oviposition period to maintain similar mite density across treatments. Eggs were left to develop for 14 days. This procedure was repeated up to generation 28.

### 4.2. Phenotypic assays

#### 4.2.1. Female longevity and fecundity assays

At generation 24, we first assayed the phenotypes of experimental females by mating them with stock males. To reduce maternal effect, we relaxed sexual selection by maintaining equal sex ratio for one generation before phenotypic assays. For longevity assays, from each replicate 30 experimental females and 30 stock males (15 fighters and 15 scramblers, non-virgin) were placed together in a common plastic container (∼3 cm^2^), with two replicate containers per line. In all fecundity and longevity assays we used virgin females, which were isolated as nymphs in separate vials. Containers were checked every other day, dead females were recorded, and the number of males was adjusted to maintain an equal sex ratio. Female longevity was analysed using a mixed-effects Cox proportional hazards model (*coxme* function in R), with line replicate included as a random effect. Survival curves for the different treatments were visualised using the *survfit* function.

Fecundity assays were performed in small glass vials (∼0.5 cm2) containing a single stock male and a single experimental female. From each line, 15 females were mated with scrambler males and 15 with fighter males. After 4 days, pairs were transferred to a fresh vial, and after an additional 4 days the mites were discarded. Eggs were counted after the first vial change and again after the second 4-day period. If a female did not lay any eggs over 8 days, we tested whether this was due to female or male sterility by pairing the female with a new male and checking for eggs after 3 days. Females that subsequently laid eggs were assumed to have been initially paired with a sterile male and their fecundity was excluded from the dataset.

Total female fecundity was modelled with stock male morph and selection regime as fixed effects. All fecundity analyses were performed using linear mixed-effects models (*lmer* function in R) assuming a Gaussian error distribution, with line replicate as a random effect. Interactions between fixed effects were initially included, but were not significant in any model and were therefore removed from the final models.

#### 4.2.2. Male harm assays

At generation F27, we quantified the effects of experimental males on the longevity and fecundity of stock females. For longevity assays, single stock virgin females were paired with either a fighter or a scrambler experimental male in individual glass vials. Males and females were isolated as juveniles and reared in glass vials until they reached maturity. From each replicate line, we tested 30 pairs in total (15 females with scrambler males and 15 with fighter males). Vials were checked every other day, and dead males were replaced with males from the corresponding experimental line.

Fecundity assays and statistical analyses were conducted as described above for the female assays.

#### 4.2.3. Inbreeding experiment

To assess inbreeding effects under different selection regimes, within each line, 16 families were established from each replicate line, with eight virgin females were individually mated with fighter sires and another eight with scrambler sires. Their progeny was isolated, and F1 females were either mated with their brothers (inbred treatment) or with a male from a different family within the same line (outbred treatment).

After egg laying, F1 adults were removed, and their progeny were placed in separate vials. From each family, a single daughter was selected and mated with a fighter male from the outbred stock population. After 4 days, the pair was transferred to a fresh vial and discarded after a further 4 days. Egg counts from the first and second vials were summed and analysed using linear mixed-effects models (*lmer*) with selection regime and treatment (inbred vs outbred) as fixed effects and line replicate as a random effect.

### 4.3. DNA isolation and sequencing

DNA samples were collected from mites at generations 1 and 28 for sequencing. To prevent contamination from food and plaster-of-Paris, the mites were transferred to dishes with a 5% agarose base and starved for three days. In total, 32 pooled samples were used for resequencing (2 generations x 4 lines x 2 treatments x 2 sexes), containing either 100 males or 100 females from each line. Males and females were selected and sequenced separately to avoid overrepresentation of DNA from considerably larger females in the pool of a given line. The sampled mites were stored at −20°C in ATL buffer until DNA extraction.

DNA was extracted using the MagJet Genomic Kit for generation 1 and the DNeasy Blood & Tissue Kit for generation 28, following the manufacturer’s protocols. The quality and quantity of the extracted DNA were assessed using Nanodrop and Qubit, respectively. Genomic library preparation was conducted using the NebNext Ultra FS II kit. Sequencing was performed at the SNP&SEQ Sequencing Facility in Uppsala on a NovaSeq 6000 system, using S4 flow cell to produce 2 x 150 bp paired-end reads.

### 4.4. Bioinformatic analyses

Reads were trimmed with Trimmomatic (v. 0.39, Bolger et al., 2014) using default settings and mapped to a reference chromosome-scale genome assembly (Chmielewski et al., 2025) using BWA-MEM (v. 0.7.17, Li, 2013). Duplicate reads and those with a mapping quality score below 20 were removed with Samtools (v. 1.9, Li et al., 2009). Quality control of the sequencing reads was conducted using FastQC (v. 0.12.1). Quality control of mapped reads was performed with qualimap (v. 2.2.1, García-Alcalde et al., 2012) and MultiQC.

Mpileup files were generated based on each of the 32 bam files using Samtools mpileup, including only positions with a base quality greater than 30. Due to decreased accuracy of SNP detection in regions flanking indels, which is caused by incorrect realignment, we removed 5 base pairs flanking indels using the filter-pileup-by-gtf.pl script. Indels were detected using the PoPoolation (v. 1.2.2, Kofler et al., 2011) script identify-genomic-indel-regions.pl with a minimum indel count of 5 and removed along with repetitive elements using filter-pileup-by-gtf.pl. The resulting mpileup file was then converted into a sync format using mpileup2sync.jar.

Our protocol was the same as in Parrett et al. (2022). Allele counts of female and male samples from the same replicate lines and time points were merged. To increase accuracy of allele frequency estimation and reduce regions with collapsed homologous sequences, we performed coverage-based filtering according to best practices for handling Pool-seq data (Kofler, Orozco-terWengel, et al., 2011). Using every 100,000th autosomal and 10,000th sex chromosome positions, we drew the distribution of the mean coverage across all samples to determine the value of the target coverage, which we define as the peak of the highest coverage density. The target coverage (101X for autosomes and 75X for the sex chromosome) was used as a reference point for filtering genomic positions. We retained only loci with mean coverage across all samples within the range of 50-200% of the target coverage (Figure S2). *R. robini* exhibits X0 sex determination system, where females carry two copies, and males one copy of X chromosome. Therefore, the difference in coverage between sex chromosome and autosomes is consistent with the expectation for a pool containing an equal ratio of sexes, as males carry a single copy of the sex chromosome, while females carry two copies of the sex chromosome (Parrett et al., 2022). The sync file, filtered for mean coverage across all lines, was split into 16 individual sex-merged lines.

### 4.5. Estimates of the autosomal genetic diversity

Because diversity estimates are sensitive to variation in sequencing coverage, we first subsampled each line to a uniform coverage of 53× using the subsample-pileup.pl script from the PoPoolation pipeline. Only autosomes were used to estimate the genetic diversity, so sex chromosome positions were excluded. We then kept only positions that met the required coverage threshold in all samples (more than 84 M positions in total). These data were used to estimate autosomal diversity patterns and to infer effective population sizes.

Genetic diversity measures, including nucleotide diversity (π) and Watterson’s estimator (θ) were calculated at both synonymous and non-synonymous sites using the Syn-nonsyn-at-position.pl script with a minimum allele count of 3 (approximately 5% of the subsampled coverage). Due to the presence of unexpectedly high number of gene models in the bulb mite gene annotation (60,309 gene models; Parrett et al. 2022), we analysed only genes showing significant expression in adult bulb mites (13,389 genes, Plesnar-Bielak et al., 2024) to avoid taking into account gene candidates with unclear functionality. Additionally, we retained only genes with coverage above 53X for at least 60% positions of gene exons (12,334 genes) in all 16 samples.

π and θ were also computed across exons using the Variance-at-position.pl script from PoPoolation software. To assess genome-wide diversity beyond genes, π and θ were estimated across the entire genome in 10kb non-overlapping windows using the script Variance-sliding.pl. Consistent with the gene-based filtering, we retained only those windows for which all samples had a minimum coverage of 53X across at least 60% of the window length.

To compare genetic diversity in all four categories (synonymous and non-synonymous positions, exons and 10kb windows), we used analysis of variance using *aov* function in R to compare the genetic diversity mean values across different experimental lines, with treatment, generation, and their interaction included as explanatory variables.

To separate the effects of selection from drift, we examined whether changes in π or θ between generations at non-synonymous sites differed between treatments while controlling for presumed neutral changes estimated from synonymous sites. Changes in non-synonymous diversity were calculated by dividing the value of π (θ) at each gene at F28 by the value at F1. For synonymous genetic variation, the genome-wide average of π (θ) at F28 was divided by the same measure at F1. To reduce data skewness and the influence of outliers, the natural logarithm of these values was calculated after adding one. The relative change in non-synonymous diversity (Δ_*π*_ or Δ_*θ*_) was calculated according to formula:

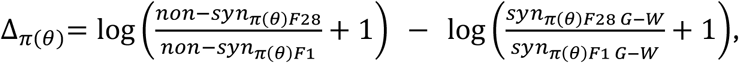

where non-syn and syn are the mean values of π or θ at non-synonymous and synonymous positions for each gene, respectively, and G-W represents the genome-wide autosomal average of π or θ. All computations were conducted using R (v. 4.1, R Core Team, 2024).

### 4.6. Effective population size estimates

Effective population size (N_e_) estimates were calculated for each line and based on allele frequency shifts between ancestral and evolved samples, separately for autosomal and sex-linked loci. The N_e_ estimates were obtained using the function estimateNe from the poolseq R package (Taus et al., 2017), with the following parameters: method = P.planII, poolSize = 200 diploid individuals, and excluding SNPs with allele frequencies below 0.01 or above 0.99.

### 4.7. Divergence between ancestral and evolved lines

To detect SNPs that changed significantly in frequency during experimental evolution, we used the ACER test (Spitzer et al., 2020), a modification of the classical Cochran-Mantel-Haenszel test, designed to detect consistent allele frequency changes across replicates. For this analysis, we used the sync file prior to coverage subsampling and further filtered SNPs to retain only those with a minor allele frequency above 5%. The test was applied using the ACER R package with the function adapted.cmh.test, run separately for male-biased and female-biased lines. Resulting *P* values were transformed to *q* values using the qvalue package (Storey et al., 2024) to control the false discovery rate. SNPs with *q* values below 0.05 were assigned to three categories: 1) diverged in both treatments, 2) diverged only in the female-biased lines and 3) diverged only in the male-biased lines. The first category contains both loci responding divergently to sex-ratio manipulation (if allele frequencies change in the opposite direction in male- and female-biased lines) and loci underlying adaptation common to the experimental conditions (if allele frequencies change in the same direction in male- and female-biased lines). To distinguish these, we estimated allele frequencies with the af() function from the poolSeq R package, averaging values within each generation and treatment. Allele frequency change for each SNP was then calculated as the difference between its frequency at generation 28 and at the starting generation. The treatment-specific categories (2 and 3) contain loci under selection in one treatment but not the other.

Due to low number of SNPs, we could not detect haplotypes which diverged between male-biased and female-biased lines. Instead, we detected haplotypes that responded exclusively in female- or male-biased lines. Haplotype identification was based on ACER test results. In E&R studies, it is commonly observed that the number of selection targets identified is much higher than expected (Kofler & Schlötterer, 2014). This is primarily due to the tight linkage between selection targets and the genetic background, which increase their frequency along with the selection target (genetic hitchhiking). Assuming that the true selection targets are among the most differentiated SNPs within each haplotype, we selected top 10% of significant SNPs with the lowest *q* values from each haplotype (Otte & Schlötterer, 2021). Haplotype identification was carried out with the haploValidate R library (Otte & Schlötterer, 2021) using mncs = 0.015 and default settings.

To make our results directly comparable to those of Parrett et al. (2022), we also performed Generalized Linear Models (GLMs) with quasi-binomial distribution to identify SNPs that consistently diverged between male- and female-biased lines during experimental evolution. The input for the GLMs was the same as that used for the ACER test but included allele counts only from F28 samples. For each genomic locus, allele frequencies were compared between the treatments using the formula: *glm(cbind(major allele count, minor allele count) ∼ treatment)*. If any line had an allele count of 0, we added 1 to the allele counts of all lines (Wiberg et al., 2017). To control for the false discovery rate due to multiple testing, *P* values obtained from the GLM were converted to *q* values as described above.

### 4.8. Gene Ontology Term Enrichment

To determine whether selection targets (defined as top 10% of significant SNPs with the lowest *q* values from each haplotype) share functional characteristics, we performed gene ontology (GO) term enrichment analysis using clusterProfiler R package. This was done separately for the SNPs that diverged in the male- and female-biased lines. GO enrichment was performed using the clusterProfiler R package (Yu et al., 2012), with Benjamini-Hochberg correction to adjust for multiple comparisons. The set of background genes comprised only genes with significant expression in adults with a high proportion of informative coverage (>60% length within 53-215X), and an assigned GO category

## 5. Data availability statement

All sequence data are available under NCBI BioProject number PRJNA1402707. Data from phenotypic assays have been published in Zenodo (https://doi.org/10.5281/zenodo.18242942) Code used in this study is available at the GitHub repository under the link: https://github.com/sebchm/sex_ratio_expEvol

## 7. Acknowledgments

We thank Małgorzata Serafin, Karolina Sobala, Józefina Wasilewska and Sylwia Jedut for laboratory assistance and Katarzyna Dudek for preparing genomic libraries. This work was supported by the National Science Centre (Narodowe Centrum Nauki) grant no. 2017/27/B/NZ8/00077 awarded to J.R. Proofreading was done with the assistance of an AI language model; the authors reviewed and approved all edits and take full responsibility for the final content.

