## Supplementary information for "Experimental evolution under biased sex ratios: phenotypic and genomic responses in the bulb mite, *Rhizoglyphus robini*"

### List of supplementary figures

**Figure S1**. Estimates of effective population sizes

**Figure S2.** Coverage distribution

**Figure S3.** Manhattan plots of SNPs which significantly diverged in both treatments between ancestral and evolved populations

**Figure S4**. Distribution of haplotype length

**Figure S5**. Allele counts from the only SNP that was diverged between treatments in the evolved lines.

### List of supplementary tables (attached separately as an Excel file)

**Table S1**. Summary of sequencing yield per sample

**Table S2**. Line-specific estimates of effective population sizes

**Table S3**. Functional description and allele frequency change in SNPs selected in opposite direction between male- and female-biased lines which were significant in both treatments.

**Table S4**. Haplotype-averaged allele frequency changes.

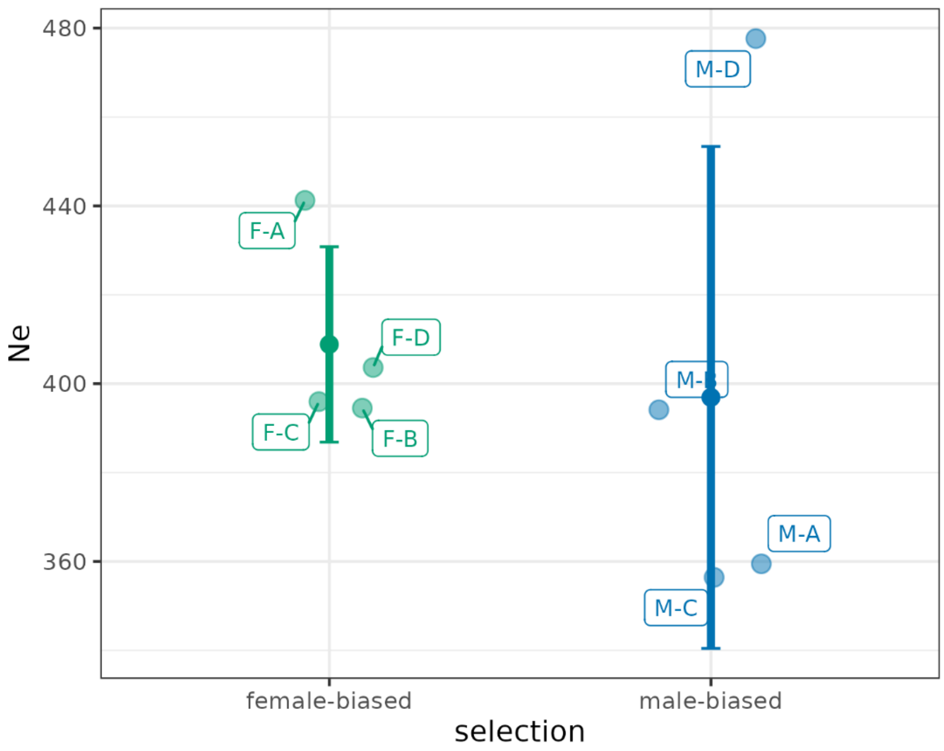


**Figure S1**. Estimates of effective population sizes based on allele frequency change between ancestral and evolved lines.

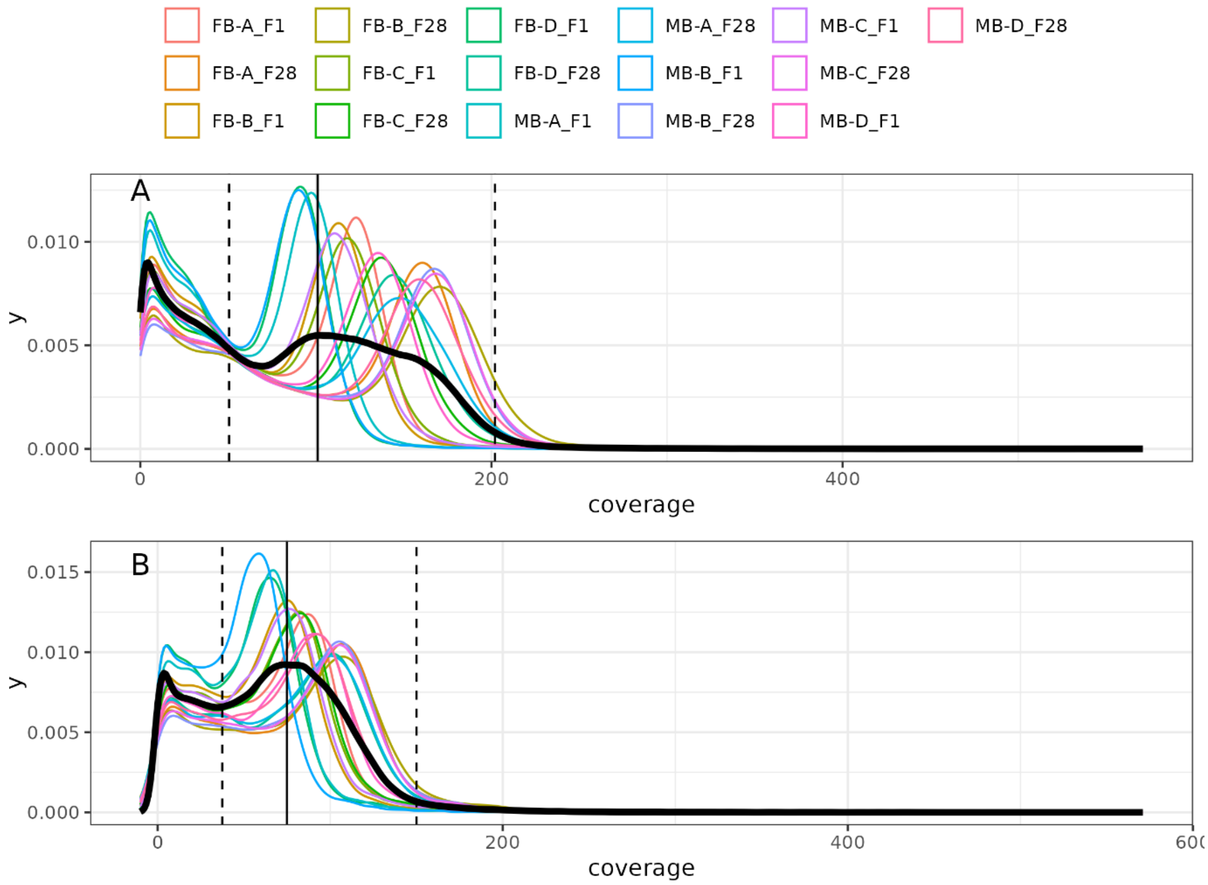


**Figure S2.** Distribution of coverage for autosomal (A) and sex chromosome (B) loci. The black smoothed line indicates the average coverage across lines. The vertical solid line represents the target coverage, while the dashed lines indicate the range of informative coverage, spanning from 50% to 200% of the target coverage. In panel A, the long tail of the coverage distribution has been trimmed to the maximum coverage value of the sex chromosome (571X), removing 12 positions with high coverage values.

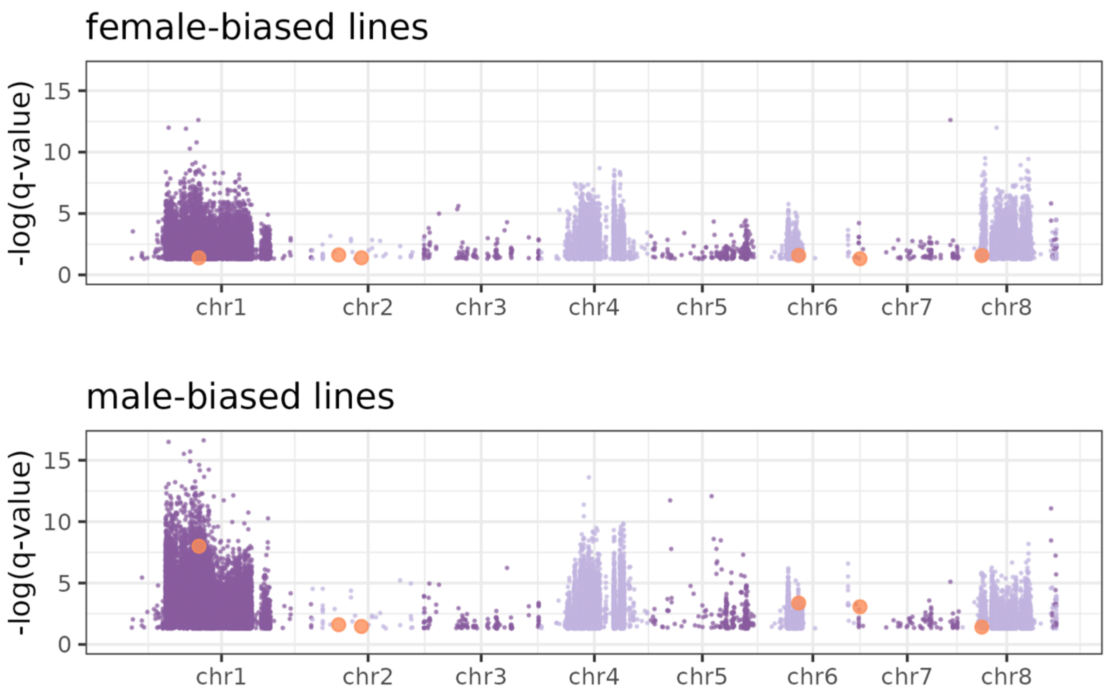


**Figure S3.** Manhattan plots of SNPs which significantly diverged in both treatments between ancestral and evolved populations. Plots show the *q* value of ACER test performed on either female-biased lines (upper panel), or male-biased lines (lower panel). All SNPs, besides these highlighted in orange, changed their frequency concordantly in both treatments. SNPs highlighted in orange changed their frequency in opposite direction between female- and male-biased lines. Colours refer to the Figure 4.

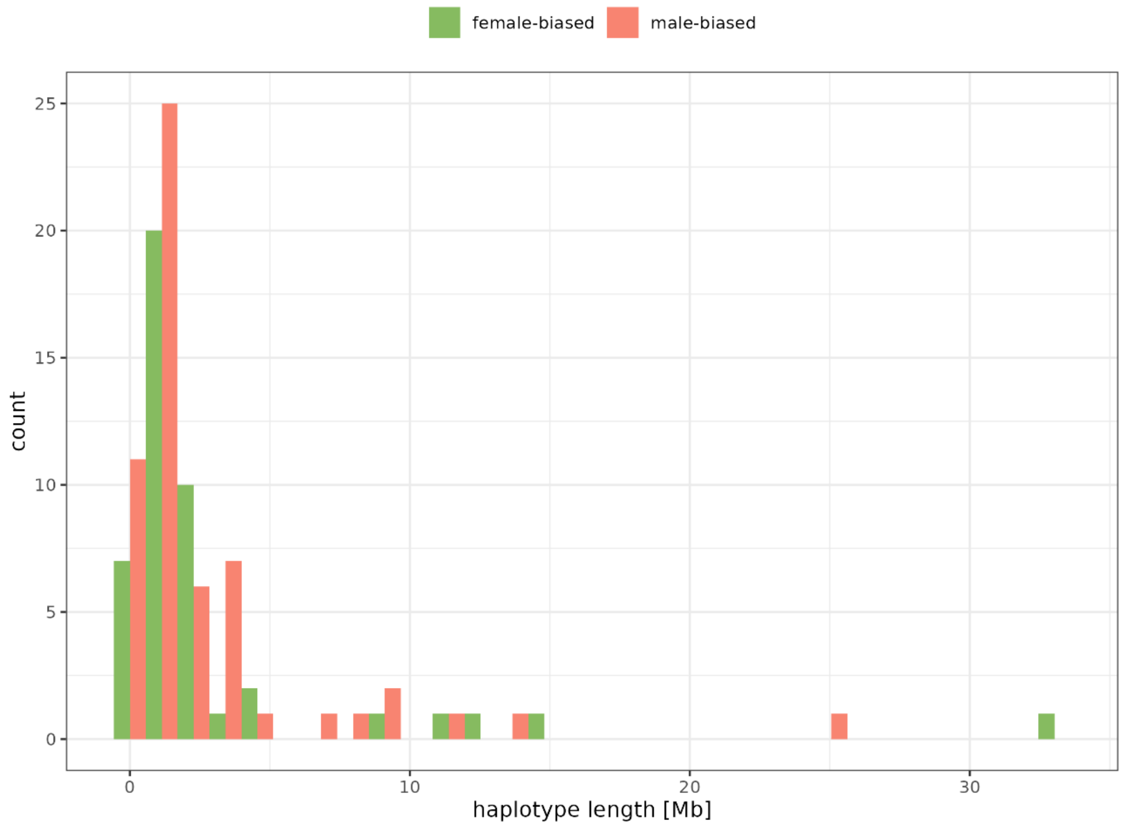


**Figure S4**. Distribution of haplotype length.

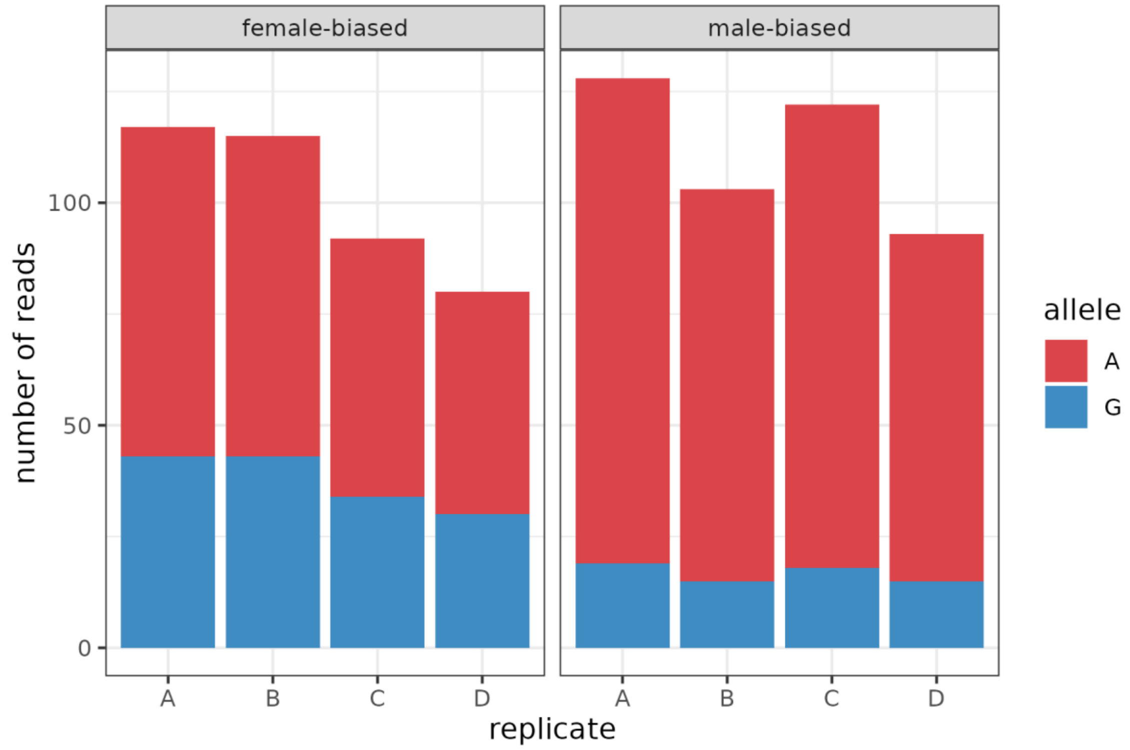


**Figure S5**. Allele counts from the only SNP that was diverged between treatments in the evolved lines, as indicated by the GLM.
